# The Evolution of Biomineral Armor in Fungus-farming Ants

**DOI:** 10.64898/2026.02.07.704540

**Authors:** Joseph A. Sardina, Gaspar Bruner-Montero, Jeffrey Sosa-Calvo, Ted R. Schultz, Hongjie Li, Cameron R. Currie

## Abstract

Biomineralization has enabled remarkable adaptations across the animal kingdom, including defensive structures and rock-grinding teeth. Although calcium carbonate is a common biogenically formed mineral found in all extant animal phyla, it is only recently known in insects and almost universally occurs with minimal magnesium content. A high-Mg calcite armor layer, covering most of the exoskeleton, was recently discovered in the fungus-farming ant *Acromyrmex echinatior*. Here, using a combination of scanning electron microscopy, energy-dispersive spectroscopy, and X-ray diffraction, paired with genome-scale phylogenetic analysis based on ultraconserved elements (UCEs), we characterize the evolution of biomineralization in fungus-farming ants. We show the presence of high-Mg calcite biomineral in 35 of 50 fungus-farming ant species studied, including taxa that span all major agricultural lineages. The biomineral is absent in all 16 myrmicine outgroup ant species, indicating an origin of the biomineral within ant agriculture, with ancestral state reconstruction analysis slightly supporting multiple origins within the fungus-farming ant clade. We then show that the occurrence of the biomineral is closely associated with the ancient mutualism with antimicrobial-producing *Pseudonocardia*, finding biomineral presence to be significantly correlated with the *Pseudonocardia* symbiont and the cuticular symbiotic structures that maintain these bacteria. Furthermore, we reveal significant inter-species variation in mineral composition and coverage, as well as intra-colony caste-specific differences, highlighting its adaptive complexity. Collectively, this work demonstrates that the innovation of a high-Mg calcite armor is strongly associated with the evolution of ant agriculture, revealing a rare biomineral to be widespread in the fungus-farming ants.

## Introduction

Biomineralization – the biological formation of minerals – has served as a key evolutionary innovation in metazoans, giving rise to structures with diverse forms, functions, and mineral compositions (1, 2). Biominerals appear in the fossil record over 540 million years ago, and likely played a role in the extreme diversification of the Cambrian explosion (3). One particularly common biomineral is calcium carbonate (CaCO_3_), which has evolved independently in numerous metazoan lineages (4). Today, calcareous biogenic structures are present in nearly all extant animal phyla (5), with examples including the hard parts of mollusk shells (6), the light-focusing eye lenses of chitons and brittlestars (7, 8), the calcareous tube-building marine serpulid worms (9), the rock-grinding teeth of sea urchins (10, 11), and the gravity-detecting crystals in vertebrate (including human) ears (12). Surprisingly, despite the diversity of the class Insecta, calcareous biominerals had not been reported from any insect until recently, with the discovery of high-Mg calcite in the fungus-farming ant *Acromyrmex echinatior* (13).

The biomineral exo-layer of *A. echinatior* is composed of ∼25-30 mol% MgCO_3_, making it one of the most Mg-enriched calcite biominerals known in nature (13). This degree of Mg enrichment is unusual, as most reported Mg calcite biominerals contain minimal amounts of Mg. The integration of Mg into calcite is not kinetically favored due to the exceptionally strong bond formed between Mg^2+^ and water (14), suggesting Mg-incorporation is highly advantageous to the ants. Mg-enrichment has been previously shown to harden and strengthen biomaterials in various organisms (15–18). Nanoindentation analysis demonstrated that the Mg calcite covered exoskeleton of *A. echinatior* ants was nearly two times as hard as the exoskeleton of mineral-free *A. echinatior* ants, despite only adding ∼7% to cuticle thickness (13). In artificial combat scenarios against *Atta cephalotes* soldier ants, which have heads twice as large and bodies ∼60% longer compared to *A. echinatior*, mineralized *A. echinatior* major workers fared significantly better than non-mineralized *A. echinatior* major workers (13). Mineralized *A. echinatior* ants lost fewer body parts, survived longer, and consistently defeated *A. cephalotes* soldiers – outcomes rarely achieved by the non-mineralized *A. echinatior* ants. Additionally, the biomineral layer was found to confer protection against entomopathogenic infection (13). Elucidating the evolutionary history and phylogenetic distribution of the biomineral in fungus-growing ants, as well as its potential association with Pseudonocardia symbionts, will provide insights into the adaptive significance of biomineralization in ants.

The fungus-farming ants (Formicidae: Myrmicinae: Attini: Attina) are a monophyletic clade of ∼250 species in 20 genera that practice insect fungiculture (19). The ants obligately cultivate a fungus within their colonies by sustaining the fungal cultivar on foraged nutrition, and subsequently consume the fungus as their primary food source (20, 21). This ancient and coevolved mutualism arose once in the Neotropics ∼66 million years ago, and has since diverged into two major clades, the Paleoattina and Neoattina, and four major agricultural systems (lower, yeast, coral-fungus, and higher agriculture) (21–23). Lower-, yeast-, and coral-fungus-farming ants forage the forest floor for detritus to incorporate into and grow their unique fungal species.

Higher fungus-farming ants grow fungi that produce specialized, nutrient-dense food bodies called gongylidia that are eaten by the ants; these ants forage for litter and debris but also collect fresh plant leaves and seedlings to feed their fungal cultivars. Leaf-cutter ants, a monophyletic group within the higher fungus-farming ants, primarily cut and harvest fresh plant material to incorporate into their fungal gardens. Resultingly, mature leaf-cutter ant colonies, which can house millions of worker ants, exhibit exceptionally high leaf-processing efficiency, making them dominant herbivores in the Neotropics (24–26). Outside the leaf-cutting ant group, fungus-farming ants form much smaller colonies, with worker numbers ranging from tens to up to a few thousand workers at a given time (27).

Fungus-farming ants have numerous defenses for protecting their resource-rich colonies, which harbor nutrient-dense ant brood and fungal cultivars. To suppress microbial threats, the majority of fungus-farming ant species maintain symbiotic *Pseudonocardia* actinobacteria on their cuticles (28). These microbial symbionts produce antibiotic compounds that inhibit the growth of garden parasites, including the specialized parasitic microfungus *Escovopsis* (29, 30). To support the *Pseudonocardia*, the ants possess specialized cuticular symbiotic structures that house and nourish the bacteria via associated gland cells (31, 32). Across the fungus-farming ant clade, symbiotic structures have convergently evolved multiple times and diversified morphologically into crypt- and tubercle-like forms. (32, 33). Within a colony, the symbiotic structures and their associated bacterial growth are typically present in female alates and workers but reduced or absent in male alates, reflecting caste-specific behaviors (33). In contrast to the males who typically do not perform colony tasks, the females indirectly apply *Pseudonocardia-*produced antibiotics to the fungus garden while tending and maintaining the garden. Moreover, the alate female reproductives vertically transmit the symbiont when founding a new colony (24, 34). In addition to garden defense, *Pseudonocardia* provides direct protection of workers against entomopathogenic fungi (35), highlighting its role as a broader defensive symbiont in this system.

Threats of larger size also regularly attack and raid the colonies, such as invertebrates, agro-predator ants, and neighboring fungus-farming ants (36–38). Non-leaf-cutting ants, which are relatively small in size and contain a single, uniform worker caste, typically rely on crypsis, death-feigning, and escaping behaviors in order to survive an attack (27, 37, 38). In contrast, leaf-cutter ant colonies, which possess a polymorphic worker caste, deploy major workers (or “soldiers”) that are up to 200 times the size of their minor workers to defend the colony (24).

Additionally, the Mg calcite biomineral of *A. echinatior* was found to serve as a defensive armor in simulated battles (13).

Given the clear functional benefits that the Mg calcite armor provides to *A. echinatior* ants, here we explore the evolutionary history of Mg calcite biomineralization across fungus-farming ants. Specifically, we examine 65 ant taxa – including all major fungus-farming ant lineages and non-fungus-farming ant outgroups – using scanning-electron microscopy (SEM), energy dispersive X-ray spectroscopy (EDS), and X-ray diffraction (XRD) analysis to detect and characterize the Mg calcite biomineral layer and estimate Mg-enrichment. We employed ultraconserved element (UCE) phylogenomics to establish a robust phylogenetic framework for tracing biomineral evolution. Ancestral state reconstruction reveals the evolutionary origins of biomineralization in the fungus-farming ants, and using this phylogenetic framework we further test for correlated evolution between the biomineral and other fungus-farming ant traits, including the *Pseudonocardia*-symbiosis. Lastly, we explore inter-caste morphological differences of the biomineral layer. By documenting the presence, composition, and evolutionary history of the Mg-enriched calcite exo-layer, this work sheds light on biomineralization as an adaptive innovation in the fungus-farming ants.

## Results and Discussion

### Widespread occurrence of a Mg-enriched calcite biomineral across the fungus-farming ants

Using a combination of SEM, EDS, and XRD (Fig. 1), we found that a Mg and Ca-enriched crystalline layer was widespread among the fungus-farming ants. Out of the 50 examined fungus-farming ant species (around one-fifth of described extant species), the biomineral was present on the worker ants of 35 (70%), and at least one mineralized taxon was found in 13 of the 19 examined genera. XRD analysis confirmed the mineral exo-layer to be a high-Mg calcite.

**Figure 1.**
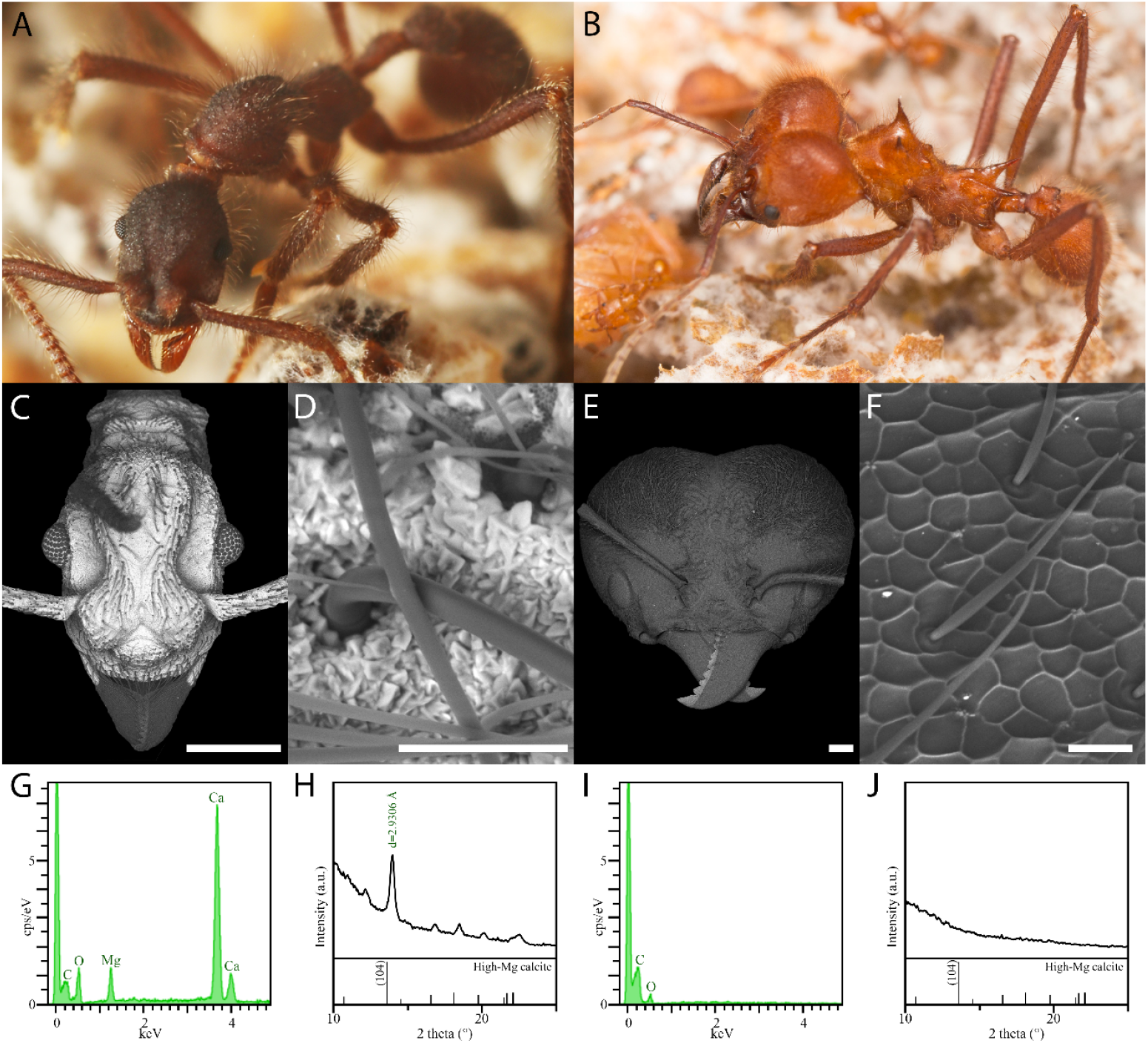
Characterization of the high-Mg calcite biomineral layer in the fungus-farming ants. (A) An *Apterostigma dentigerum* worker ant. (B) Extremely large caste size polymorphism between a minor worker (bottom left corner) and a super soldier from the same *Atta cephalotes* colony. (C-F) SEM-BSE images of *Apterostigma auriculatum* (C, D) and *Atta cephalotes* (E, F) reveal presence and absence, respectively, of the crystalline exo-layer. (G) SEM-EDS analysis of the *Apterostigma auriculatum* mineral layer showing elemental composition (C, O, Mg and Ca). (H) XRD analysis confirms the Mg- and Ca-bearing exo-layer of *Apterostigma auriculatum* as a high-Mg calcite. High-Mg calcite reference pattern shown at bottom. (I) SEM-EDS analysis of the *Atta cephalotes* cuticle. (J) XRD analysis of *Atta cephalotes*, indicating no crystalline structure. Images (A) and (B) captured by Don Parsons, and used with permission. Scale bars: 500 µm (C,E); 25 µm (D,F).

Morphologically, the biomineral was most commonly found covering the antennal scape, head, thorax, petiole, legs, and abdomen of the worker ants, with some intriguing exceptions to this pattern described below (Fig. 2, right panel). Interestingly, all mineralized ants lacked the mineral around cuticular joints and on the sensory organ-dense antennal funiculi; we suspect that mineralization of the cuticle in these regions would result in suboptimal functionalization (e.g., inhibiting range of motion, and dampening external sensory detection). Of note, mineralized taxa spanned both major sister clades of the fungus-farming ants, the Paleoattina and Neoattina, as well as the four major agricultural groups: lower agriculture, yeast agriculture, coral fungus agriculture, and higher agriculture (including leaf-cutting ants).

**Figure 2.**
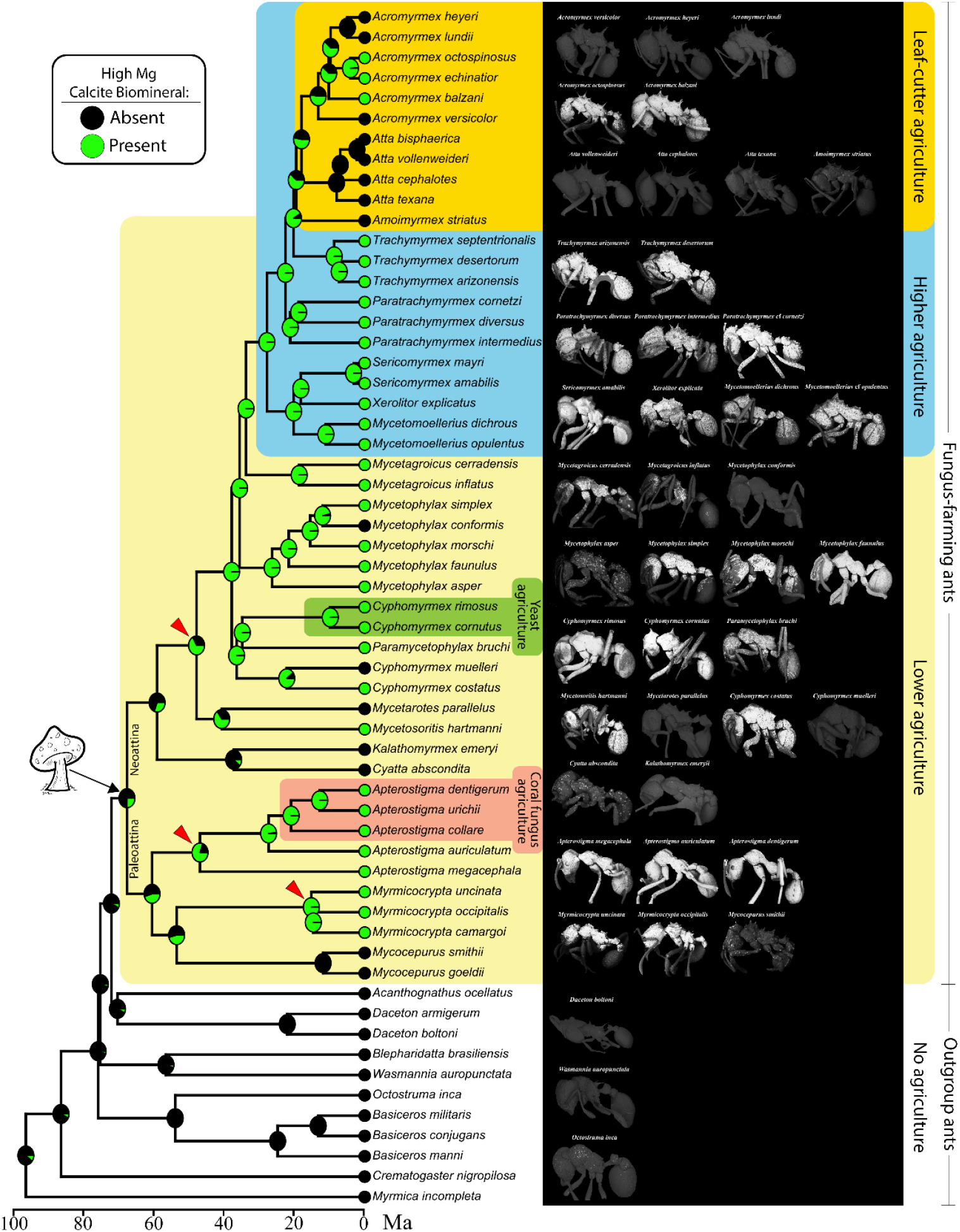
The high-Mg calcite biomineral is widespread across the fungus-farming ants. Left: A time-calibrated phylogeny of the fungus-farming ants based on a UCE phylogenomic analysis indicating presence (green) or absence (black) of the high-Mg calcite exo-layer. Probabilities from ancestral-state reconstruction analysis are indicated by the circles at the ancestral nodes, which suggest three likely origins of the biomineral within the Attina (red arrowheads). Right: Whole body SEM-BSE images demonstrating the coverage of the Mg calcite layer in the mineralized taxa. Mushroom pictograph drawn by Claudia Kepler.

Within the Paleoattina clade, eight out of ten examined species possessed the biomineral layer. The only Paleoattina genus in which no biomineral was found was *Mycocepurus* (*M. goeldii* and *M. smithii*). In contrast, all five examined species of *Apterostigma*, including both coral and lower fungus-farming species, had a uniform crystalline layer covering nearly their entire external surface, including the antennal scape, head, thorax, petiole, abdomen, legs, as well as the crevices between individual ocelli of the eyes. The crystal morphology in most of the *Apterostigma* taxa (*A. auriculatum, A. collare, A. dentigerum*, and *A. urichii*) was similar, possessing large and bulky crystals (∼ 5-7 µm in diameter). However, the crystal morphology in *A. megacephala*, the earliest-diverging *Apterostigma* species, was flat and relatively smooth such that individual crystals were difficult to identify. Lastly, within the *Myrmicocrypta* clade, all species examined (*M. camargoi, M. occipitalis*, and *M. uncinata*) possessed the crystalline layer. In *M. occipitalis* and *M. uncinata* the mineral covered the head (including eye crevices), thorax, and petiole, but was absent from the antennae, legs, and abdomen. The complete lack of mineral on the abdomen makes the body coverage pattern of these *Myrmicocrypta* taxa one of the most unique amongst the fungus-farming ants and suggests tight regulatory control of biomineral formation across the body. Given this striking contrast of biomineralized and biomineral-free cuticle on a single ant, *Myrmicocrypta* may serve as a useful model in future studies aiming to interrogate the biomineralization process and the underlying genetic regulation.

In the Neoattina clade, which diverged from the Paleoattina over 60 million years ago (Fig. S5)(21), the biomineral was detected in all three major agricultural groups – lower agriculture, yeast agriculture, and higher agriculture (including leaf-cutter ants). The two species belonging to the earliest-diverging lineage of the lower Neoattina, *Cyatta abscondita* and *Kalathomyrmex emeryi*, lacked the biomineral layer. Of note, SEM-BSE detected extraneous masses of heavy elements sparsely populating the cuticle on *C. abscondita*, but SEM-EDS confirmed these to be soil debris (Fig. S6). Two taxa representing the next earliest diverging Neoattina lineage differed in biomineral presence – *Mycetosoritis hartmanni* was covered in a mineral layer and *Mycetarotes parallelus* exhibited a smooth, biomineral-free cuticle. Out of five taxa examined in the *Cyphomyrmex*-*Paramycetophylax* clade, four possessed an external mineral layer (*Cyphomyrmex cornutus, Cyphomyrmex costatus, Cyphomyrmex rimosus*, and *Paramycetophylax bruchi*), with the exception of the mineral-free *Cyphomyrmex muelleri*. The head and dorsal abdomen of *C. rimosus* possessed large regions of low biomineral abundance. In *P. bruchi*, weakly biomineralized regions also regularly populated the external surface of the head and thorax, and the abdomen was mostly bare of the biomineral except for the most posterior regions. However, in *C. cornutus* and *C. costatus* the mineral layer abundantly covered the head, thorax, petiole, and abdomen. Of note, the biomineral layer on *C. cornutus* formed around the symbiotic crypt structures, such that the entrances to these invaginated crypts remain open, and the crystals surrounding the crypts are larger in size relative to the crystals not surrounding the crypts (Fig. S7). In five *Mycetophylax* species examined, the crystalline layer was detected in four taxa, but the amount of mineral varied greatly among the species. *Mycetophylax faunulus* was nearly completely covered in a biomineral layer, whereas *Mycetophylax simplex* and *Mycetophylax morschi* had the mineral present on most of the head, thorax, and abdomen but with some bare or spotty regions. Biomineral was detected on *Mycetophylax asper*, but only in small, regular patches across the body. No mineral was detected on *Mycetophylax conformis*. Lastly, both taxa representing the sister lineage to the higher fungus-farming ant clade, *Mycetagroicus inflatus* and *Mycetagroicus cerradensis*, possessed the mineral. However, both taxa had a high proportion of their surface lacking biomineralization relative to most other mineralized ants. The reduced amount of biomineral abundance seen in *Mycetagroicus* and most species of *Mycetophylax* may be due to biological effects of the specimens analyzed, such as a younger age of the ants (prior to complete biomineralization) or a colony-specific nutritional deficit. However, this coverage pattern appears to be correlated with phylogeny, which suggests that this may not be an anomaly of sample size or collection bias. Whether or not this reduced coverage pattern is a result of relaxed selective pressure in these lineages to maintain the mineral layer is an interesting future question.

All 12 non-leaf-cutting, higher fungus-farming ant species examined were covered in the biomineral layer. *Mycetomoellerius dichrous* and *Mycetomoellerius opulentus* had areas of large mineral crystals interspersed with regions of thin mineral layers which were in between, but not covering, the cuticular papillae (Fig. S7). The bodies of *Sericomyrmex amabilis, Sericomyrmex mayri*, and *Xerolitor explicata* were nearly completely covered in large crystals, covering all external regions minus the antennal funiculus, mandibles, and ocelli. All three examined *Paratrachymyrmex* species (*P. cornetzi, P. diversus*, and *P. intermedius*) were also nearly completely covered in the mineral layer, although *P. diversus* had regular, small mineral-free regions (∼6 µm diameter) likely corresponding to the underlying symbiotic structures (Fig. S7). Similarly, all four examined *Trachymyrmex* species examined (*T. arizonensis, T. desertorum, T. saussurei*, and *T. septentrionalis*) were mineralized, with *T. arizonensis, T. desertorum*, and *T. saussurei* possessing a mineral layer that covered most of the cuticle excluding around the areas with tubercular symbiotic structures. It is intriguing that all of the higher, non-leaf cutter species examined here possessed a mineral layer. This could be the result of strong selective pressure that has been maintained on these taxa due to a unique aspect of higher fungus-farming ant lifestyle, such as: predation pressure, defensive behavior, nest architecture (e.g., gas exchange), diet, or colony size.

Surprisingly, the leaf-cutters were the group with the most biomineral-free taxa examined in this study. No mineral was found in *Amoimyrmex striatus*, which represents the earliest-diverging leaf-cutter lineage. All *Atta* taxa studied (*At. bisphaerica, At. cephalotes, Atta texana*, and *At. vollenweirdi*) had smooth, mineral-free cuticles, consistent with their shiny cuticles observable to the naked eye. On the other hand, a mineralized exo-layer was detected on four of seven *Acromyrmex* species. The earliest-diverging *Acromyrmex* species, *Ac. versicolor*, as well as *Ac. heyeri* and *Ac. lundii* had cuticles similar to *Atta* – smooth and mineral-free. However, *Ac. balzani, Ac. echinatior, Ac. octospinosus*, and *Ac. volcanus* all possessed the mineral layer. The coverage of the mineral layer was similar in *Ac. echinatior, Ac. octospinosus*, and *Ac. volcanus*, with head and thorax covered in biomineral, and posterior abdomen, antennae, and legs weakly covered or absent; *Ac. volcanus* had spotty coverage on the thorax as well. *Ac. balzani* had a unique mineral coverage amongst the mineralized *Acromyrmex* species, with the head, thorax, petiole, abdomen, and legs completely covered in the mineral layer. The only fungus-farming ant genus not examined in this study was *Pseudoatta*, a monotypic genus which is morphologically similar to, and phylogenetically nested within, the *Acromyrmex* clade. Given that this genus is a known social parasite to *Acromyrmex lundii* and *Acromyrmex heyeri* – two species found here lacking the biomineral – it is unlikely that *Pseudoatta* possesses a mineral layer, although future studies should confirm this. Nonetheless, the absence of the mineral layer in many leaf-cutter ant species is interesting. The leaf-cutters engage in the most derived form of agriculture, in which they cut and process fresh vegetation to incorporate into their fungal cultivars. This highly-efficient farming results in extremely large mega-colonies, which can produce millions of worker ants and high numbers of reproductive male and female alates. Other aspects of the leaf-cutter colonies, for example the size and power of the *Atta* soldier ant caste and the unusually large number of worker ants, may have relaxed selective pressure to maintain the protective biomineral armor.

### Mg calcite biomineral not found outside of the fungus-farming ants

To determine if the Mg calcite layer is present outside the fungus-farming ant group, we next examined 15 outgroup ant species. Of these, seven (*Acanthognathus ocellatus, Blepharidatta brasiliensis, Crematogaster nigropilosa, Daceton armigerum, Daceton boltoni, Myrmica incompleta*, and *Wasmannia auropunctata*) had smooth external surfaces characteristic of a typical ant cuticle. The other eight species, *Basiceros conjugans, Basiceros manni, Basiceros militaris, Eurhopalothrix bruchi, Octostruma inca, Pogonomyrmex pima, Strumigenys gundlachi*, and *Talaridris mandibularis*, were examined due to dull (non-shiny) cuticles observed from light micrographs; these taxa did not have smooth cuticular surfaces and instead had an accumulation of extraneous matter covering sparse regions of the exoskeleton. However, XRD analysis of these ants did not detect a Mg calcite signal, and EDS revealed these areas to be composed of mostly C, O, S, Si, and Fe, and sometimes Al (Fig. S6). Given the abundance of iron-oxide and quartz (silicon-dioxide) in soil, these EDS results are consistent with reports of the accumulation of soil debris on the exoskeleton of these taxa (39).

Taken together, we did not detect any Mg calcite crystalline biomineral outside of the Attina. Although our sampling of non-fungus-farming ants was necessarily non-exhaustive due to the species richness of Formicidae and the throughput capabilities of XRD, our results are consistent with a recent large-scale study employing X-ray microtomography, which identified a conspicuous, highly X-ray-absorbing cuticular layer in multiple fungus-farming ant species but did not detect comparable structures in any non-fungus-farming ant (40). X-ray microtomography does not resolve mineral phase or composition, and therefore cannot distinguish biomineral crystallization from other heavy-element-containing materials; however, the absence of a highly X-ray-absorbing cuticular layer in non-fungus-farming ants provides strong evidence against the presence of a dense mineralized coating outside of the fungus-farming ants. The results reported here, together with a lack of reported Mg calcite in the widely diverse insect class, suggest that the Mg calcite biomineral exo-layer is a relatively unique innovation of the fungus-farming ants.

### Evolutionary history of Mg calcite biomineral in the fungus-farming ants

To investigate the evolutionary history of the Mg calcite biomineral in fungus-farming ants, we next performed maximum-likelihood ancestral state reconstruction analysis. The results indicated that the last common ancestor of the fungus-farming ants did not possess the Mg calcite layer, and that there were three separate subsequent origins of the biomineral – two in the Paleoattina clade and one in the Neoattina clade (Fig. 2, left panel). Incorporating the node dating estimates of our phylogenetic tree, we then deduced the timing of these origins. The single origin within the Neoattina clade was estimated to be the earliest of the three origins, dating back to ∼ 47.97 mya. The second origin, at the last common node of *Apterostigma* in our phylogeny, was estimated at ∼46.98 mya. The third estimated origin was relatively recent, at the last common node of *Myrmicocrypta* in our phylogeny, dating back to ∼21.8 mya. Interestingly, these three origins closely coincided with the previously reported three independent origins of the fungus-farming ant symbiotic structures (33). Further, the two oldest estimated origin events, dated around 48 and 47 million years ago, were at a time when atmospheric carbon dioxide levels were extremely high. A recent study estimated CO_2_ concentrations peaked around 50 mya at about 1600 ppm, almost four times higher than current levels (41).

Support for these inferred origins is modest at several key nodes, and alternative evolutionary scenarios remain plausible. Alternatively, given the uncertainties and assumptions in ancestral state and transition rate estimates, the biomineral may have originated a single time near the origin of fungus-farming ant agriculture, followed by multiple subsequent loss in descendant lineages. Thus, while our results marginally support multiple origins of Mg-calcite biomineralization, they also point to a complex evolutionary history. Future analyses incorporating fossil-calibrated data, for example XRD analyses of fungus-farming ants preserved in amber, could help further resolve the timing and history of biomineral evolution in the fungus-farming ant clade.

### Correlated evolution between the biomineral and *Pseudonocardia*, the symbiotic structures, and papillae

Given the similar origins of the biomineral (this study) and the fungus-farming ant symbiotic structure (see (33)), we next conducted correlated evolution analyses of the biomineral versus three different characters: (i) association with *Pseudonocardia* bacteria, (ii) presence of symbiotic structures, and (iii) presence of cuticular papillae structures. All three characters were found to have interdependent evolutionary histories with the biomineral, as indicated by significantly high Akaike Information Criterion (AIC) weights (0.999983, 0.999994, and 0.999954, respectively; Fig. 3). Given these results, and that the majority of mineralized species also possess all three of the other traits, this supports a possible functional and developmental connection between these fungus-farming ant traits and biomineralization.

**Figure 3.**
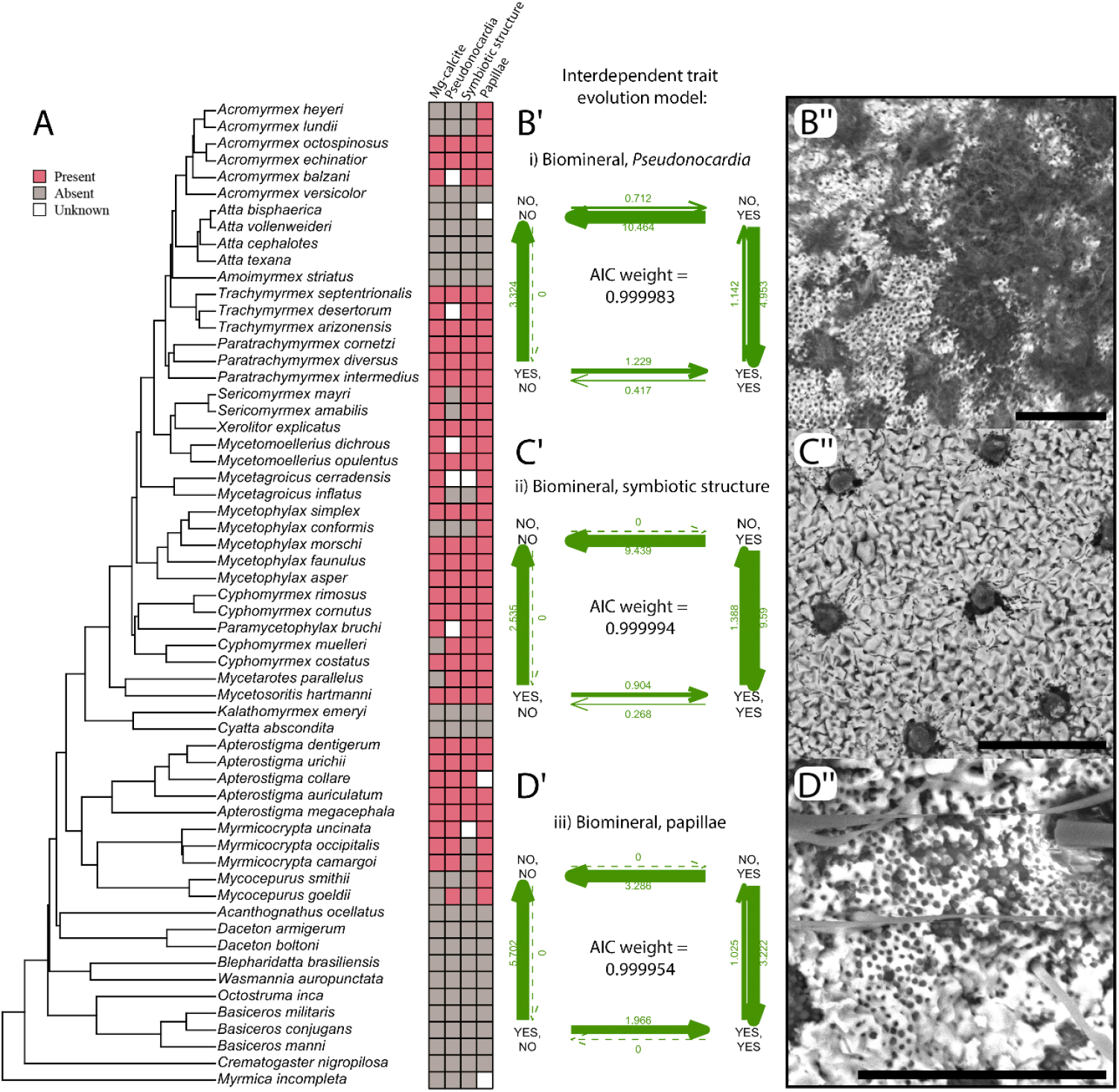
Correlated evolutionary analysis of the Mg calcite biomineral and three traits of interest: (i) association with *Pseudonocardia* bacteria, (ii) presence of symbiotic structure, and (iii) presence of papillae. (A) Phylogeny of the fungus-farming ants and outgroups with boxes on the right indicating presence (red), absence (gray), or unknown state (white) of the biomineral and the three traits of interest. (B’, C’, D’) Interdependent trait evolution model of the biomineral and three ant traits of interest. In each panel, trait states represent combinations of presence (‘YES’) or absence (‘NO’) of the traits indicated (e.g., “YES, NO” represents biomineral presence and second trait absence). Arrows illustrate estimated transition rates between states, with thickness proportional to transition rate magnitude and numbers indicating rate estimates. AIC weights of the respective analysis indicated in the center. (B”, C”, D”) SEM-BSE images showing the biomineral with the respective trait of interest. Scale bars = 30 µm.

Out of 28 mineralized taxa with available data, 25 of them have been reported to maintain a symbiosis with *Pseudonocardia*, with the exception of *Mycetagroicus inflatus, Sericomyrmex amabilis*, and *Sericomyrmex mayri*. In contrast, *Cyphomyrmex muelleri, Mycetarotes parallelus*, and *Mycocepurus goeldii* have been reported with *Pseudonocardia* growth on their exoskeleton but were found here with no mineral layer. Interestingly, we also found filamentous bacteria on the abdomen of *Myrmicocrypta*, despite having a completely biomineral-free abdomen (Fig. S8). Whether *Pseudonocardia* plays a role in the formation of the high-Mg calcite ant biomineral layer remains unknown. However, the capacity of some autotrophic Actinobacteria to sequester environmental CO_2_ has been documented (42, 43), suggesting a potential role of the ant symbiont in a CO_2_-fixation process underlying carbonate formation on the ant exoskeleton.

Further, our SEM imaging revealed distinct filamentous-shaped depressions on the external surface of the biomineral layer (Figure S9, also see examples in Figure S8), resembling “micro-fossils” that are produced by calcite-depositing marine bacteria likely by the bacterial cell surface and associated extracellular polymeric substances acting as mineralization nucleation sites 44– 46). Though the similarities are intriguing, whether the *Pseudonocardia* symbiont is performing these biomineralization-related activities on the ant cuticle remains to be tested. Notably, consistent with the exceptions noted above, mineralized *Sericomyrmex* and *Mycetagroicus* that lack *Pseudonocardia* associations provide valuable comparative cases for future work.

The symbiotic structure-biomineral interdependence had the largest AIC weight out of the three analyses, concordant with the fact that most mineralized species also possessed symbiotic structures (28 out of 31 taxa with available data); however, there were taxa that only possessed one character and not the other. For example, *Mycetagroicus inflatus, Myrmicocrypta occipitalis*, and *Myrmicocrypta uncinata* form the biomineral but have no symbiotic structures present, whereas *Cyphomyrmex muelleri* and *Mycetarotes parallelus* possess crypt- and tubercle-shaped symbiotic structures, respectively, but were found lacking a mineral layer. It has been previously reported that the majority of fungus-farming ants possess these specialized cuticular symbiotic structures, and their presence is highly correlated in extant taxa with the presence of the *Pseudonocardia* bacteria (33). This is plausible considering their current known function of housing and feeding the exoskeleton-dwelling *Pseudonocardia*. Whether the symbiotic structure plays a role in the formation of the biomineral remains to be tested. One hypothesis is that the symbiotic structures directly secrete molecules required for Mg calcite production. Previous work has identified a gland cell connected to the symbiotic structures (32), which has been hypothesized to secrete nutrients for *Pseudonocardia*; however, the chemical composition of the secretion is unknown. On the other hand, if the *Pseudonocardia* is directly aiding in biomineral formation, the symbiotic structure may contribute by mediating *Pseudonocardia* growth. Of note, the mineralized *Sericomyrmex* taxa possess symbiotic structures despite lacking an association with *Pseudonocardia* (33); this may suggest, at least in the *Sericomyrmex* lineage, an alternative, non-symbiosis-related function of the symbiotic structures.

Lastly, all mineralized fungus-farming ant species with available data also possessed cuticular papillae (32/32 taxa with available data). Papillae are small cuticular structures roughly one micron in diameter and height, although these structures may have slight inter-species variation. On the fungus-farming ants with papillae present, these structures are extremely abundant, populating nearly the entire surface of the cuticle; for example, we found *Acromyrmex echinatior* major worker ants to have roughly 350,000 papillae per mm^2^. Interestingly, in *Myrmicocrypta* the papillae are present on the head, thorax, and petiole, but are completely lacking on the abdomen, closely resembling the body coverage of the biomineral in *Myrmicocrypta*. Currently, there is no known function of the papillae. From SEMs, we have observed that the biomineral appears to form in between and on top of the papillae. One possible function may be biomineral attachment, in which the knob-like papillae structures help fix the Mg calcite layer to the ant cuticle. Alternatively, the papillae may secrete organic compounds that aid in the biomineral formation. These hypotheses remain to be tested. Of note, seven extant species possessed papillae but lacked a mineral layer (*Acromyrmex heyeri, Acromyrmex lundii, Mycetophylax conformis, Cyphomyrmex muelleri, Mycetarotes parallelus, Mycocepurus smithii*, and *Mycocepurus goeldii*). Nonetheless, the fungus-farming ant papillae have received virtually no scientific investigation, therefore, due to the strong correlation found here between papillae and biomineralization, these small but peculiar structures warrant future investigation.

### Variation in Mg calcite biomineral composition across the fungus-farming ants

To characterize differences in biomineral composition, we compared XRD patterns, specifically the value of the d104 peak, which decreases due to the smaller size of Mg atoms relative to Ca. These results revealed large interspecific variation, with a maximum d104 peak of 3.0127 Å (*Paramycetophylax bruchi*) and a minimum of 2.9221 Å (*Sericomyrmex amabilis*) (Fig. 4). We then estimated Mg-enrichment (mol% MgCO_3_) of the biomineral based on the d104 value according to previously established methods (47). These d104-based calculations suggest considerable differences in biomineral Mg content, with estimates ranging from 8.0 to 51.3 mol% MgCO_3_. Further analysis, including using quantitative electron probe micro-analysis, should confirm these estimates. Nonetheless, the estimated Mg content present within the biomineral layer for all examined ant species is above the typical ∼4 mol% threshold that relates to changes in calcite structure and stability, above which Mg substitution significantly affects crystal properties and solubility. Thus, species found here with extremely high estimated Mg-enrichment may be of interest for future nano- and bio-material-focused investigations, given the high kinetic energy required to incorporate Mg^2-^ into the calcite crystalline structure. Additionally, we found that (104) peak d-values often varied across body regions of individual ants. These patterns typically held for multiple ants of the same species and colony, suggesting that this variation is likely biologically stable. For example, the majority of *Apterostigma* species examined had the highest estimated mol% Mg in the abdomen relative to the head and thorax; differential transcriptomic or proteomic analysis across body regions may inform on the molecules and mechanisms underlying such high Mg-enrichment.

**Figure 4.**
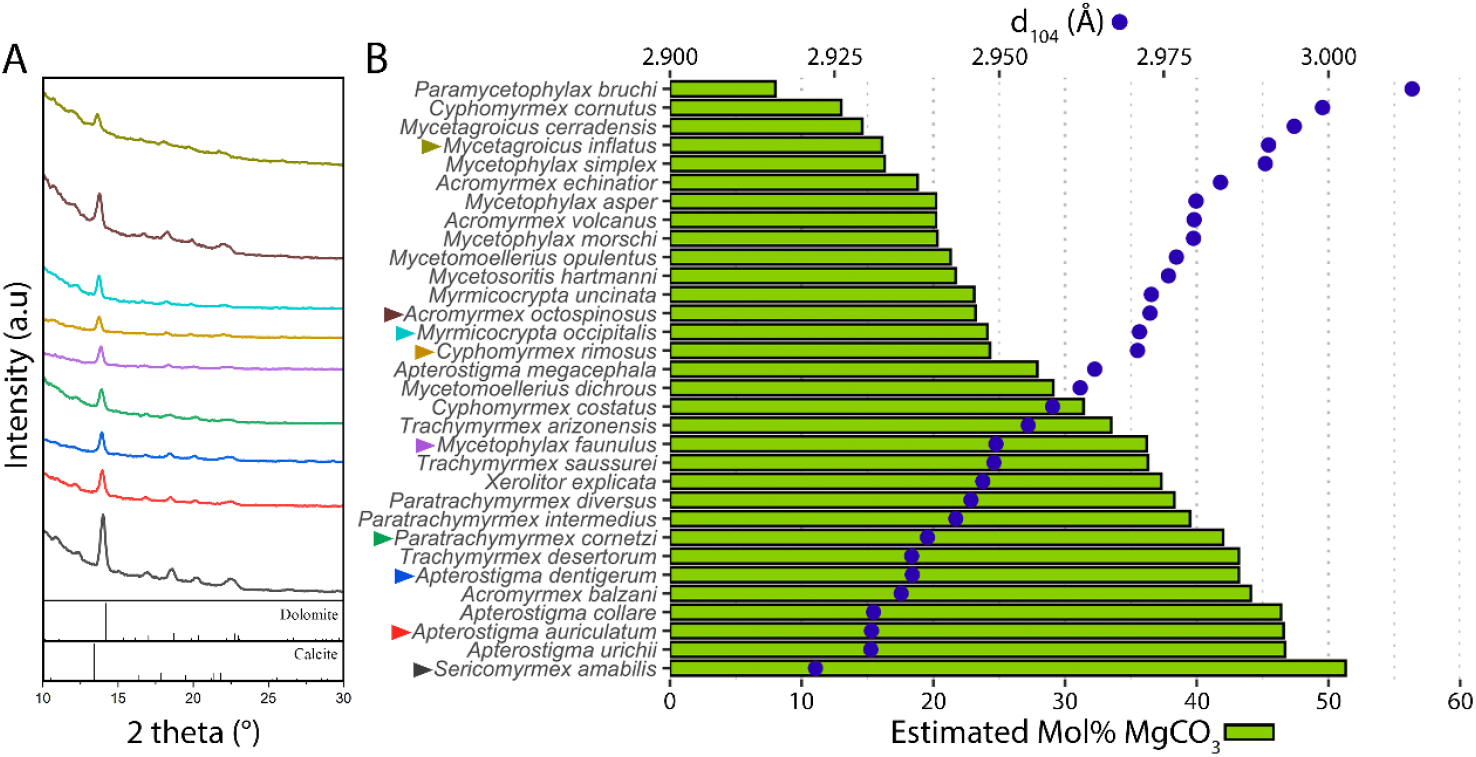
Variation in biomineral composition across mineralized fungus-farming ant taxa. (A) XRD patterns of select fungus-farming ant species (arrowheads in B). Reference patterns of dolomite (CaMg(CO_3_)_2_) and calcite (Ca(CO_3_)_2_) shown at the bottom. (B) For each mineralized taxon, the d104 peak from XRD analysis is plotted in indigo on the upper X-axis, and the estimated mol% MgCO_3_ is plotted in green on the lower X-axis.

### Caste polymorphic differences in biomineralization

We next looked for biomineral differences between castes from the same colony (Fig. 5). Ants typically have a minimum of three castes within a single colony – a reproductive male caste, a reproductive female caste, and a non-reproductive female worker caste. The worker caste can be further divided into polymorphic sub-castes; however, across the fungus-farming ants, distinct polymorphic worker sub-castes are only present in the leaf-cutter ants. We focused on four taxa representing four major fungus-farming ant systems. In the coral fungus-farming *Apterostigma dentigerum*, all three castes had a mineral layer covering the cuticle of the head, thorax, abdomen, legs, and antennal scape (Fig. S10). Two minor differences in mineral coverage were observed, in which the *A. dentigerum* females, but not the males, had mineralization in the crevices between individual ocelli and the posterior corner of the external mandibular margin. The lower fungus-farming *Cyphomyrmex costatus* had more pronounced differences between male and female castes. In this species, both sexes had mineralization on the head, thorax, abdomen, and the outer legs; however, the female worker and reproductive castes had visibly more abundant mineral coverage across the entire body relative to the male (Fig. S10). In the higher fungus-farming *Sericomyrmex amabilis*, the male reproductive caste completely lacked a biomineral layer, whereas the female worker and reproductive castes both had abundant biomineralization covering the antennal scape head, thorax, abdomen, and legs (Fig. S10).

**Figure 5.**
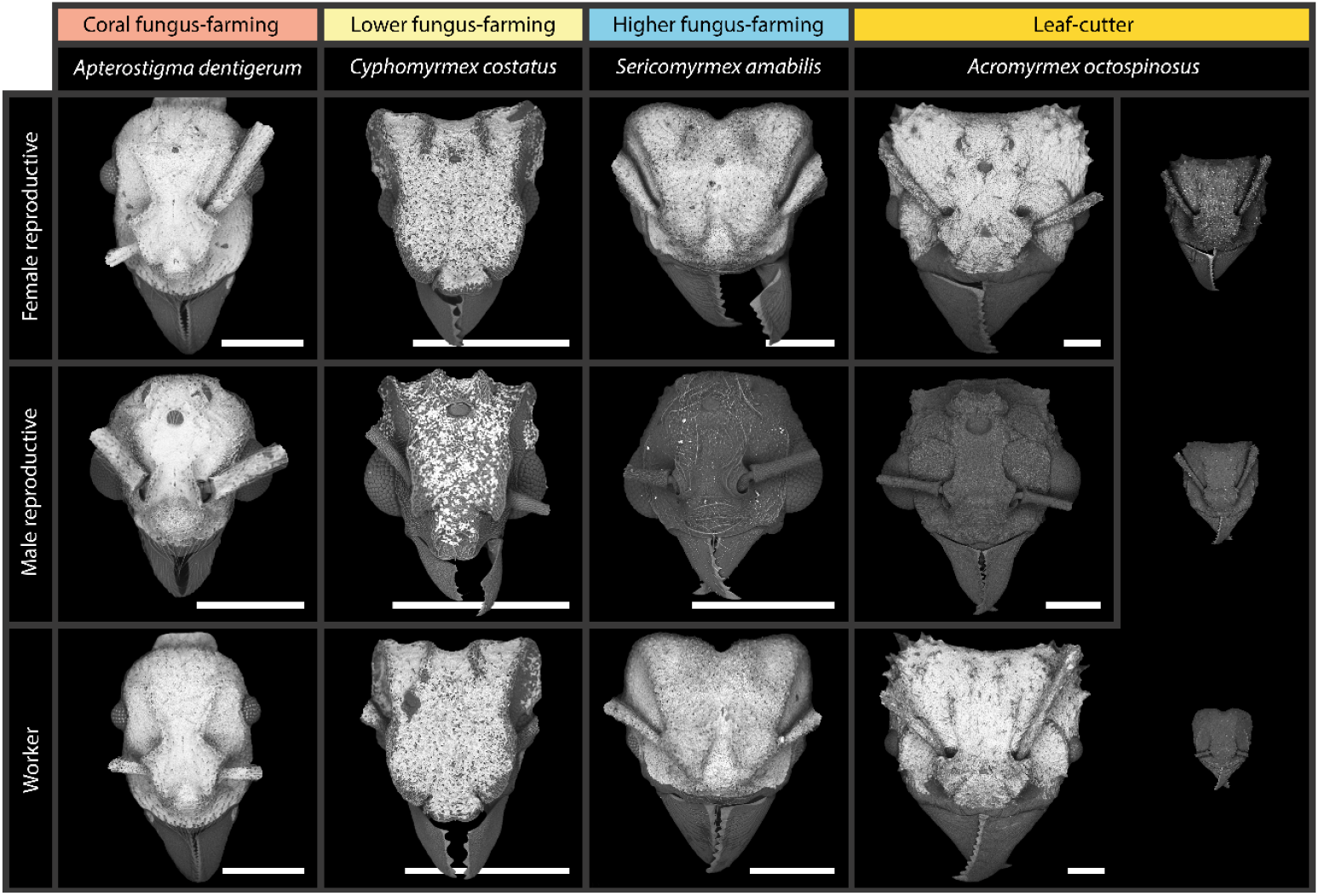
Biomineralization varies across intra-colony castes. SEM-BSE images of ant heads from each caste of four species representing four major agricultural groups: coral fungus-farming ants; lower fungus-farming ants; higher fungus-farming ants; and leaf-cutter ants. Leaf-cutter ant colonies display distinct polymorphic worker sub-castes, whereas the fungus-farming ants possess a monomorphic worker caste. Scale bar for all images is 500 µm.

Similarly, the reproductive male of the leaf-cutter *Acromyrmex octospinosus* lacked a biomineral layer, and the reproductive female ant had full biomineral coverage on the antennal scape, head, thorax, abdomen, and legs (Fig. S11). Intriguingly, we also observed differences between the worker sub-castes of *A. octospinosus* (Fig. S11). The major worker had a full mineral layer similar to the female reproductive, although the mineral on the abdomen and legs were less abundant on the major worker. The larger media worker had sparse coverage (∼5-30 microns in size) of Mg- and Ca-containing crystalline layers on the cuticle, though due to the low abundance of these areas these could not be confirmed as Mg calcite with XRD. In contrast, the mineral was completely absent from the smaller media and minor workers.

The differences in biomineral formation across castes make it a unique example of a caste-level polymorphic trait other than size or behavior. Given the high-energetic cost likely required to produce the biomineral armor, this intra-colony variation may reflect higher caste-level colony dynamics. For example, male alates typically do not perform colony-related duties and instead leave the colony shortly after eclosion to attempt to inseminate a female, and die shortly after whether successful or not. Thus, selective pressure to maintain a mineralized male ant has likely been relaxed, resulting in the mineral-free males. Therefore, the discovery of the fully mineralized *A. dentigerum* male raises the speculation of a unique lifestyle for this caste of this species, potentially including male-male combat, colony task responsibility, colony defense, or higher risk of entomopathogenic infection. On the other hand, both female castes, reproductive and worker, exhibit similar biomineralized layers, likely reflecting the similar roles carried out by these ants. Upon successful fertilization, the female reproductive ant will attempt to found her new colony, and, due to the initial lack of workers, will perform all necessary colony tasks, including foraging for fungal substrate. However, after her first brood of workers emerges and take over colony tasks, the queen’s role shifts almost exclusively to egg-laying. The biomineralization variation observed in the polymorphic worker castes of the leaf-cutter *A. octospinosus* may further reflect worker subcaste-specific roles. For example, the defensive function of biomineral armor is likely to aid the major workers, who are susceptible to attacks during foraging and colony defense, more than the minor workers, who largely remain inside the colony and have a minimal role in colony defense; these behavioral differences may also expose the major workers to entomopathogenic infection, which the biomineral layer confers protection against (13). Together, these observations suggest biomineralization is finely tuned to the specific colony roles of each caste, highlighting the interplay between structural innovation and complex social organization.

## Conclusion

In this study, we show that high-Mg calcite biomineralization is widespread across the fungus-farming ants and has a complex evolutionary history closely associated with the origin and diversification of ant agriculture. Rather than arising as an isolated novelty, our findings suggest that biomineralization in this system is consistently coupled with a broader defensive strategy that integrates microbial association, specialized cuticular architecture, and mineralized armor. The coincidence of geologic periods of exceptionally high atmospheric CO_2_ with the inferred timing of biomineral origins raises the possibility that global carbon chemistry during the early diversification of fungus-farming ants influenced both the feasibility and adaptive value of carbonate-based biomineralization. At finer biological scales, the pronounced interspecific, anatomical, and caste-specific variation we document suggests that biomineralization is dynamically regulated across colony roles, underscoring how evolutionary innovations in eusocial organisms can be differently deployed within colonies rather than uniformly manifested at the species level. Together, these results broaden the evolutionary context of Mg-enriched calcite biomineralization beyond the marine environment and show that many fungus-farming ants produce a biologically rare mineral, while exhibiting extensive natural variation across lineages, body regions, and castes.

## Materials and Methods

### Ant Sampling and biomineral detection

We examined 65 ant taxa, including 50 fungus-farming ant species spanning all major agricultural groups and 15 non-fungus-farming outgroup species, for the presence or absence of a cuticular biomineral layer. Biomineralization presence or absence was scored from scanning electron microscopy (SEM) with back-scatter electron (BSE) detector imaging, energy-dispersive X-ray spectroscopy (EDS) mapping, and confirmed by in situ X-ray diffraction (XRD) to identify mineral phase and estimate Mg-enrichment (mol% MgCO_3_). Specimen metadata (Table S1) and a detailed explanation of analytical procedures are provided in the SI Appendix.

### Molecular phylogenetic framework

DNA shearing, library preparation, and library pooling followed previously reported methods (19, 48). For enrichment of UCE loci, we used myBaits (Arbor Biosciences, Ann Arbor, MI, USA) UCE Hymenoptera 2.5Kv2A kit targeting 2524 conserved loci (49). We inferred maximum likelihood trees using IQ-TREE (50) from a 166-taxon concatenated UCE dataset under alternative filtering schemes (SI Appendix, Methods; Figs. S1-S4), and nodal support was quantified using 1,000 ultrafast bootstrap replicates. Divergence times were estimated with MCMCTREE (51) (SI Appendix, Methods; Figs. S5), and the resulting time-calibrated phylogeny was used for downstream analysis. Full details of UCE data generation and processing, and phylogenetic and divergence analyses are provided in the SI Appendix.

### Ancestral State Reconstruction and Correlated Evolutionary Analysis

Ancestral state reconstruction (ASR) and correlated evolutionary analyses were performed in R v. 4.3.3 using R packages *ape* v. 5.7.1 (52) and *phytools* v. 2.1.1 (53). A one-parameter (“equal rates”) maximum-likelihood ASR was run using the time-calibrated phylogeny (Fig. S5) which was pruned to remove taxa without biomineral presence/absence data available, and the taxa with biomineral data were coded as (0) absent or (1) present for the biomineral layer. Origin age estimates were based on the median age of the nodes from the unpruned tree (Fig. S5). The same time-calibrated phylogeny (Fig. S5) was pruned and used to test for correlated evolution between biomineral and three traits: (i) *Pseudonocardia* association, (ii) specialized cuticular symbiotic structures, and (iii) cuticular papillae structures; for each analysis, only taxa with data for both traits were included, and model support was evaluated using Akaike Information Criterion (AIC) weights.

## Acknowledgments

We thank Bil Schneider for expert assistance in SEM work; Yihang Feng, Noah Brown, and Tianyu Zhou for XRD guidance; Eugenia Okonski (National Museum of Natural History) for help with specimen and data processing; and the Wisconsin Insect Research Collection and Charlotte Francouer for the loan of ant specimens. This study was funded by National Institute of Food and Agriculture, United States Department of Agriculture grant ID number 1023233 (to C.R.C.), and the National Science Foundation grant DEB-1927155 (to C.R.C.); G.B.-M. was supported by the National Secretariat for Science and Technology of Panama (SENACYT) under grant FID24-073. We thank the Ministry of Environment of Panama (MiAMBIENTE) for granting the scientific collecting permit ARBG-050-2023, which made the fieldwork possible.

